# Antibody-Based Affinity Cryo-Electron Microscopy at 2.6 Å Resolution

**DOI:** 10.1101/064188

**Authors:** Guimei Yu, Kunpeng Li, Pengwei Huang, Xi Jiang, Wen Jiang

**Affiliations:** Markey Center for Structural Biology, Department of Biological Science, Purdue University, West Lafayette, IN, USA; Divisions of Infectious Diseases, Cincinnati Children's Hospital Medical Center, Cincinnati, OH, USA; Department of Pediatrics, University of Cincinnati College of Medicine, Cincinnati, OH, USA

**Keywords:** antibody-based affinity grid, affinity cryo-electron microscopy, single particle 3D reconstruction, Tulane Virus

## Abstract

The affinity cryo-electron microscopy (cryo-EM) approach has been explored in recent years to simplify and improve the sample preparation for cryo-EM. Despite the demonstrated successes for low-concentration and unpurified specimens, the lack of near-atomic structures using this approach has led to a common perception of affinity cryo-EM as a niche technique incapable of reaching high resolutions. Here, we report a ~2.6 Å structure solved using the antibody-based affinity grid approach with a Tulane virus sample of low concentration. This is the first near-atomic structure solved using the affinity cryo-EM approach. Quantitative analyses of the structure indicate data and reconstruction quality comparable to conventional grid preparation method using samples at high concentration. With the shifting of bottlenecks of cryo-EM structural studies to sample grid preparation, our demonstration of the sub-3 Å capability of affinity cryo-EM approach indicates its potential in revolutionizing cryo-EM sample preparation for a broader spectrum of specimens.

## Introduction

The remarkably improved image quality provided by direct electron detectors, together with other hardware and software advances, have resulted in an explosion of near-atomic resolution structures determined by single particle cryo-EM (Bai et al., 2015; Banerjee et al., 2016; Bartesaghi et al., 2015; Campbell et al., 2015; Grant and Grigorieff, 2015; Merk et al., 2016; Wang et al., 2014). Furthermore, the recent breakthrough in the Volta phase plate technique (Danev and Baumeister, 2016; Khoshouei et al., 2016) holds great promise to extend single particle cryo-EM to more samples with molecular masses below 100 KDa. Since microscopes, detectors, and image processing algorithms are no longer bottlenecks, the major hurdles for most cryo-EM projects have now shifted to sample grid preparation, which usually consists of a multistep sample purification process and the subsequent preparation of a thin film of frozen-hydrated sample on TEM grids (Grassucci et al., 2007). Further innovation in cryo-EM sample grid preparation is of great significance for developing single particle cryo-EM into a routine structural biology tool. The affinity cryo-EM approach that modifies TEM grids with an extra affinity layer to immobilize, purify, and concentrate target samples possesses a great potential to simplify and improve the cryo-EM grid preparation for a broader range of specimens, such as those of low yield and unpurified samples (Glaeser, 2015; Taylor and Glaeser, 2008; Yu et al., 2016a).

Multiple affinity cryo-EM approaches based on different types of affinity layers including functionalized lipid layer (Azubel et al., 2004; Benjamin et al., 2016; Kelly et al., 2008; Medalia et al., 2002), 2D crystals of streptavidin (Han et al., 2012), antibody layer (Yu et al., 2014) and chemically functionalized carbon surface (Llaguno et al., 2014) have been reported. The affinity cryo-EM method provides various advantages. First, it enables single particle cryo-EM studies of low-concentration samples that are often encountered due to low abundance in natural sources, low yield of expression systems, or safety concerns for highly contagious/dangerous pathogens. Secondly, it allows direct isolation of target particles from crude extracts via a specific interaction (Kelly et al., 2008; Yu et al., 2014), which combines sample purification with grid setup, simplifying cryo-EM grid preparation into a single-step process. Moreover, better sample integrity can be obtained for labile macromolecular complexes with affinity cryo-EM approaches by avoiding multiple biochemical purification steps. Finally, immobilization of particles to an affinity layer can reduce particle diffusion and minimize potential sample damages at the air-water interface during and after sample blotting (Taylor and Glaeser, 2008). Altogether, these benefits of affinity cryo-EM will potentially bring more samples within reach of single particle cryo-EM, and also provide a more convenient, higher throughput, and mild cryo-EM grid preparation method.

Although multiple 3-D reconstructions have been reported with affinity cryo-EM approaches (Han et al., 2012; Kelly et al., 2008; Llaguno et al., 2014; Yu et al., 2014; Zhang et al., 2015), these structures were all limited to low to intermediate resolutions and no near-atomic reconstructions have been obtained. Thereby there has been a common worry about using affinity cryo-EM approach for high-resolution single particle reconstructions, which prohibits the widespread use of the affinity cryo-EM approach. Tulane virus (TV) is a monkey calicivirus in the *recovirus* genus of the *caliciviridae* and serves as a surrogate for human norovirus studies (Wei et al., 2008; Yu et al., 2013). Due to the low yield, the purified TV particles from cell cultures were often at a concentration of ~10^10-11^ particles/ml, which is too low for conventional cryo-EM studies that usually need ~10^14^ particles/ml or higher for most viral samples. In this work, we have used this challenging low-concentration TV sample to explore the potential of using antibody-based affinity grid for high-resolution cryo-EM. By concentrating TV particle via the anti-TV antibody affinity layer on the grid, and also optimizing the cryo-EM grid freezing condition to achieve an optimal freezing where particles of interest immobilized on the antibody-affinity layer were frozen in vitreous ice with a thickness slightly larger than the particle size, we have obtained high quality cryo-EM data for TV particles. A ~2.6 Å structure of TV was solved using the antibody-based affinity cryo-EM approach, which for the first time demonstrated the ability of affinity cryo-EM approach for solving sub-3 Å structures.

## Results and Discussion

TEM grids were coated with anti-TV IgGs aided by protein A as described previously (Yu et al., 2016a, 2014). As expected, TV particles were immobilized and concentrated efficiently on the anti-TV antibody-coated TEM grid to a level allowing for cryo-EM grid preparation (figure s1). Cryo-EM sample of TV on the antibody-coated affinity grid was prepared using perforated carbon grids layered with thin continuous carbon (Figure 1A). The extra affinity layer brings more background noise to cryo-EM images relative to standard cryo-EM grids, which is essentially the major reason for the common concern about applying affinity cryo-EM for high-resolution cryo-EM reconstruction. The extra noises from the affinity layer cannot be computationally removed in most affinity cryo-EM approaches (Kelly et al., 2008; Llaguno et al., 2014; Yu et al., 2014). However, such loss of image contrast due to the affinity layer can be compensated by reducing the ice thickness without exposing the immobilized particles to the air-water interface. For example, reducing ice thickness by ~30 nm could theoretically balance out the extra electron scattering by an affinity layer formed by protein A and IgGs (~13 nm in thickness) on the carbon film (~3 nm) (Vulović et al., 2013). Therefore, to pursue the best possible affinity cryo-EM images, freezing conditions were screened for an optimal ice thickness that is slightly larger than the size of TV particle (~40 nm).

**Figure 1.**
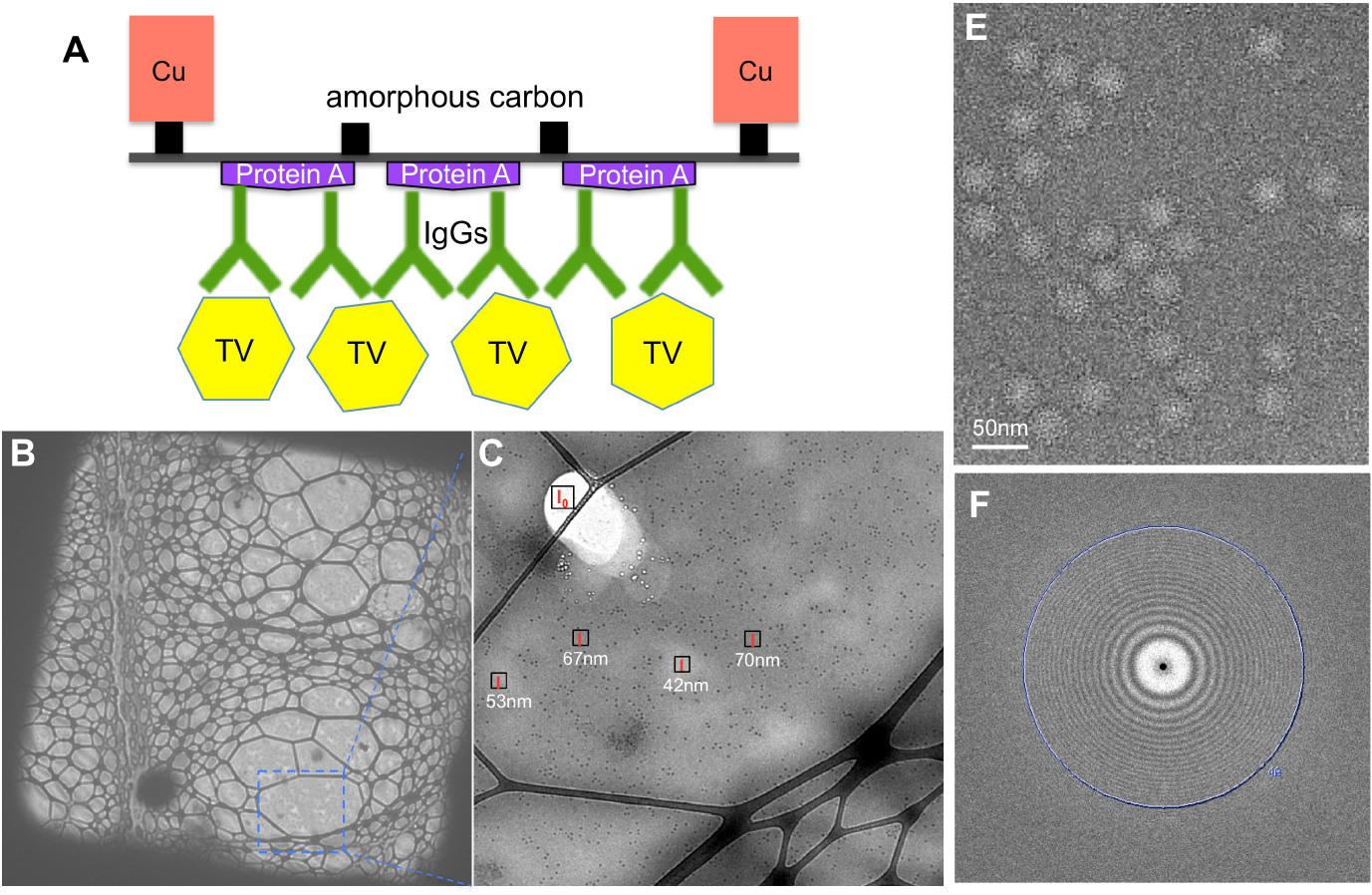
Antibody-based affinity cryo-EM of TV. (A) Schematic diagram of TV immobilized on the antibody-based affinity grid. For clear visualization, it was not drawn to scale. (B-C) Affinity cryo-EM micrographs of TV frozen using the optimized condition at 500× (B) and 3,800× (C) magnifications. Local ice thicknesses were estimated using the Beer-Lambert law based method. (E) A representative motion-corrected micrograph. (F) The 2-D power spectrum of the micrograph in (E). The position of 4 Å is labeled.

Since visual evaluation of ice thickness in the range of tens to hundreds of nanometers (e.g., 50-150 nm) is challenging, quantitative estimation of ice thickness is necessary. An ice thickness measurement method based on the Beer-Lambert law was adopted (Yan et al., 2015), in which the ice thickness (t) can be approximated from the ratio of image intensity of a melted area (I_0_) and an area with ice (I), and the inelastic mean free path of electrons in vitreous ice (λ_in_) using the equation 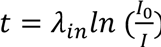. Mean image intensities (I_0_ and I) can be measured directly using the DigitalMicrograph software (Gatan, Inc.) controlling the camera, which allows convenient online measurement of ice thickness during sample screening. The quantitative estimation of ice thicknesses allowed successful optimization of ice thickness with obviously improved image contrast (figure s2). A sample freezing condition producing vitreous ice with an estimated thickness of ~40-70 nm was obtained for TV particles on the anti-TV antibody-coated grid (Figure 1B,C). Cryo-EM movies of TV were collected automatically by using Leginon (Carragher et al., 2000) to control the FEI Titan Krios microscope and the Gatan K2 Summit direct electron detector. For most movies, signals at 4 Å or higher resolutions were recovered after motion correction (Figure 1E,F).

Single particle analysis was performed using *jspr* program (Guo and Jiang, 2014). The truly independent image processing strategy was adopted, in which the whole dataset was split into two halves and processed completely independently (Guo and Jiang, 2014; Liu et al., 2016). Based on the 0.143 cutoff criterion of gold-standard Fourier shell correlation (FSC) curve, a final 3-D reconstruction of TV at an overall resolution of 2.6 Å was obtained from ~14K particles (Figure 2A,B). As shown in Figure 2A, TV structure contains two major density layers, the basal shell layer (green-colored) and the protruding layer (blue-colored). Conformational flexibility has been suggested for the protruding layer of TV and TV-related structures (Chen et al., 2006; Katpally et al., 2010; Prasad, 1999; Yu et al., 2013). To evaluate the resolutions of the two layers, FSC curves were calculated for the shell layer and the protruding layer, respectively. As expected, the shell layer with an estimated resolution of ~2.5 Å was better resolved than the more flexible protruding layer (~2.7 Å) (Figure 2B). Analysis of the determined Euler angles of TV particles revealed a uniform coverage of the angle space (figure s3), suggesting that no view preference was introduced by the antibody affinity layer.

**Figure 2.**
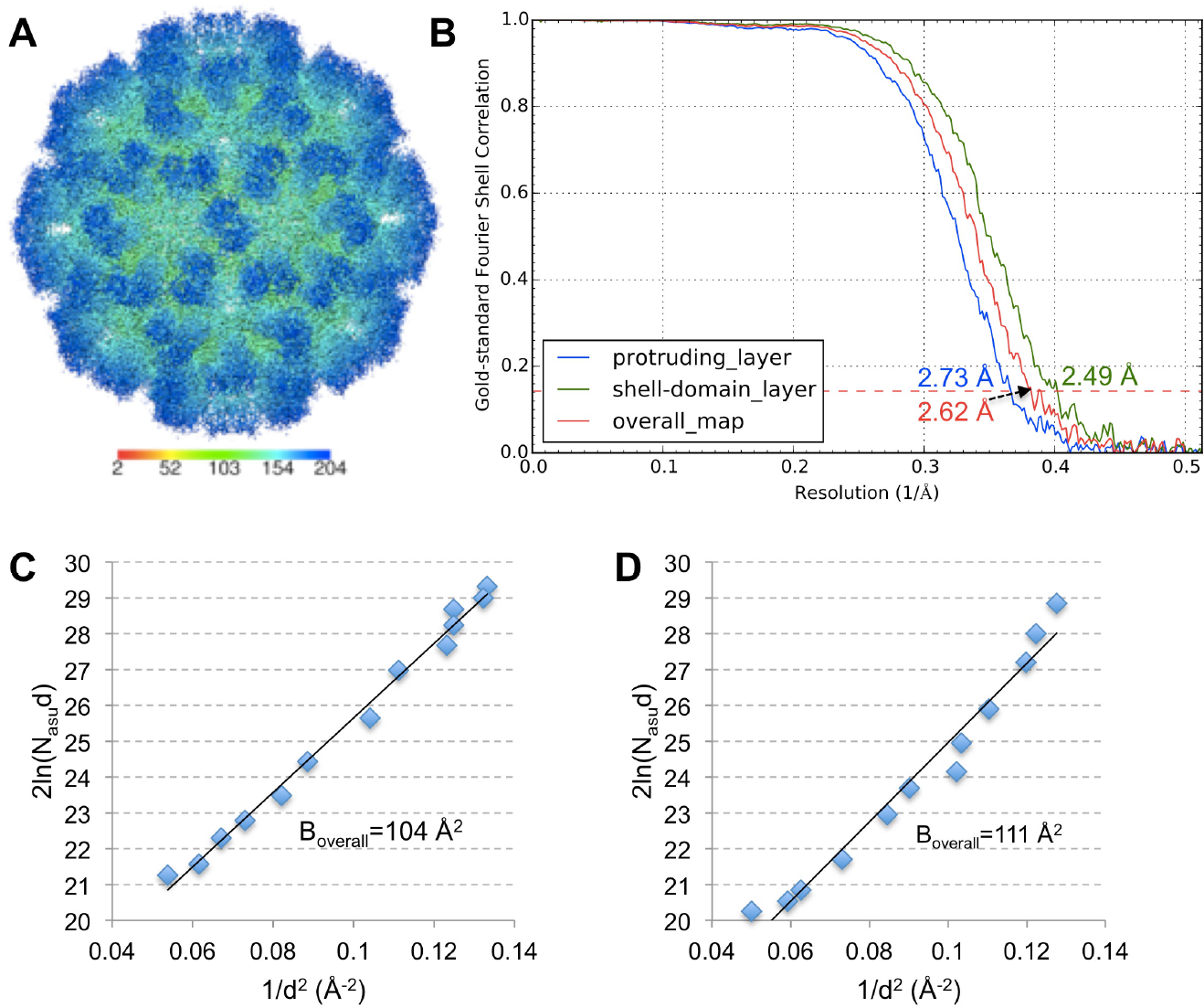
Single particle 3-D reconstruction of TV using affinity cryo-EM data. (A) Surface representation of TV radially colored according to the provided color key. (B) Gold-standard FSC curves of the overall map (red), the protruding layer (blue) and the shell layer (green). The position of the 0.143 FSC cutoff for resolution estimation is labeled with a dashed line. (C, D) *2ln(N_asu_d)* vs *1/d^2^* is plotted for *B_overall_* estimation of the affinity cryo-EM data of TV (C) and the cryo-EM dataset of 20S proteasome (EMPIAR-10025) re-processed using *jspr* (D), respectively.

To further evaluate the quality of this affinity cryo-EM TV dataset and 3-D reconstruction, its overall B factor (*B_overall_*) was approximated. *B_overall_*, which reflects the composite effect of sample integrity and homogeneity, ice thickness, imaging quality and computing errors, is a good indicator of cryo-EM data and 3-D reconstruction quality (Liu et al., 2007; Rosenthal and Richard, 2003). Based on the linear relationship between *2ln(N_asu_d)* and *1/d^2^*, in which *N_asu_* is the number of asymmetric units used for the 3-D reconstruction and *d* is the resolution, *B_overall_* is equal to the slope of the linear fitting (Rosenthal and Richard, 2003). As shown in Figure 2C, the *2ln(N_asud_)-1/d^2^* plot of this affinity cryo-EM TV dataset showed an apparent linear relationship and yielded a *B_overall_* of 104 Å^2^. This *B_overall_* is smaller than previously reported *B_overall_* for structures determined at 3.2 to 7 Å resolutions (Danev and Baumeister, 2016; Liu et al., 2007; Wang et al., 2014), which further confirms the excellent overall quality of this TV dataset. Moreover, we reprocessed a published cryo-EM dataset of 20S proteasome (Campbell et al., 2015) in the same way as for TV using *jspr* and obtained a 3-D reconstruction of ~2.7 Å (figure s4). The 20S proteasome data was collected under similar imaging conditions as TV (i.e., Titan Krios and Gatan K2 Summit) but using standard holey grids and samples. The plot of *2ln(N_asu_d)-1/d^2^* for the 20S proteasome yielded a *B_overall_* of 111 Å (Figure 2D) that is comparable to the *B_overall_* of the affinity cryo-EM TV dataset. This comparison suggests that the antibody-based affinity grid approach can not only provide cryo-EM grids of similar quality (if not better) as the standard cryo-EM grid preparation approach, but also provide distinct advantages for those samples beyond reach of standard single particle cryo-EM.

With our ~2.6 Å 3-D reconstruction of TV, many high-resolution structural features were resolved (Figure 3). Figure 3 shows three representative regions of TV density map including a short α-helix followed by a loop (Figure 3A) and two β-strands (Figure 3B,C), in which side chain densities can be easily recognized for most residues. Both hydroxyl groups in Tyr residues and the backbone carbonyl oxygen atoms are distinguishable (Figure 3A to C). In addition, this high-quality map of TV also allowed unambiguous assignment of rotameric conformations of most side chains. For example, different rotamers of Ile residues were unambiguously identified based on the density (Figure 3D) and the correct rotamers were distinguishable from multiple similar rotameric states (Figure 3E). Many ordered water molecules were also observed to form hydrogen bonds with neighboring residues or other water molecules (Figure 3F).

**Figure 3.**
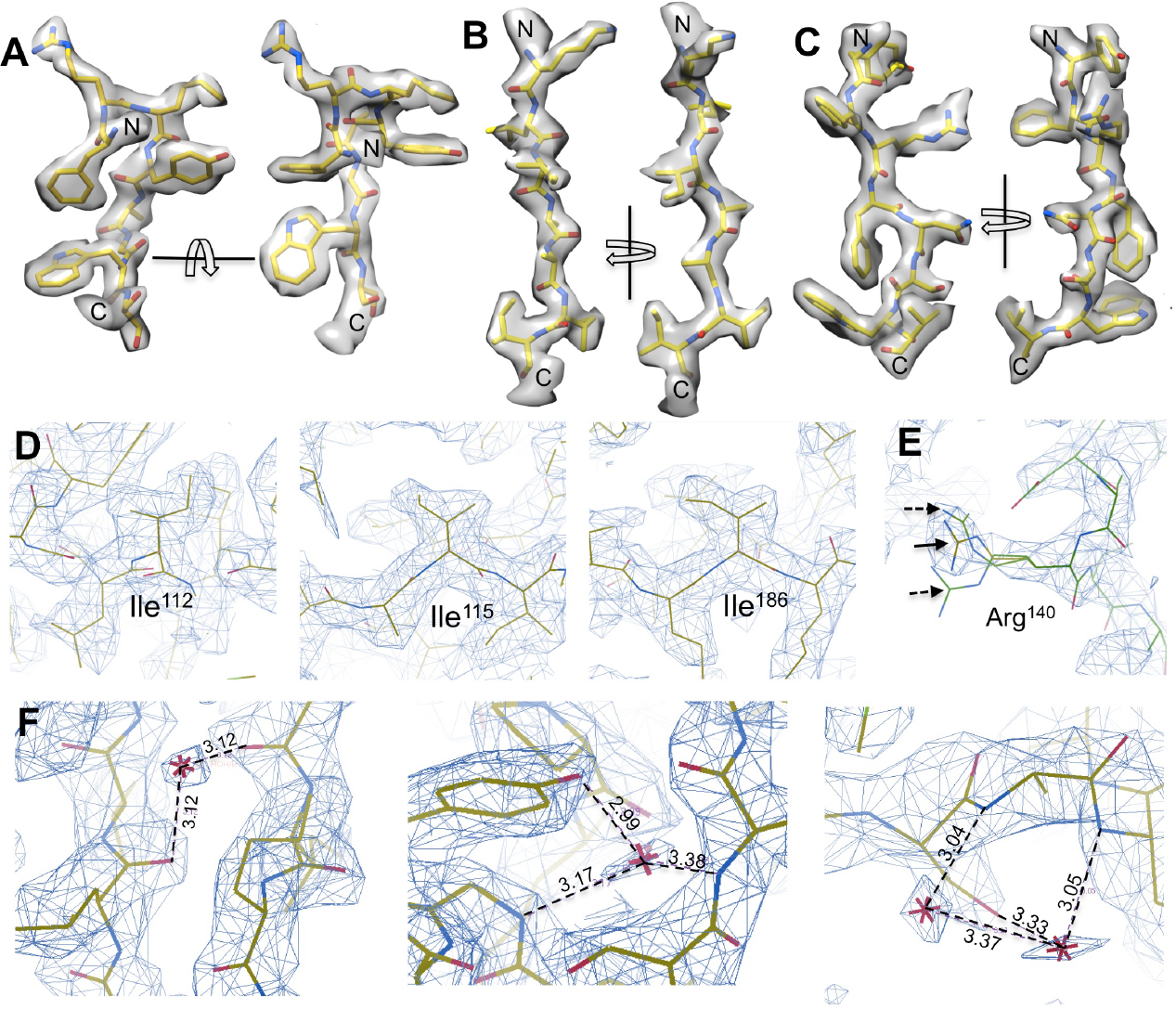
Atomic structural features resolved in the ~2.6 Å map of TV. (A) Densities for a short α-helix followed by ashort loop. (B, C) Densities for two β-strands. (D) Multiple rotameric states identified for Ile residues. (E) shows one example of distinguishing the correct rotamer (solid arrow) from alternative rotameric states (dashed arrow) for residue Arg^140^. (F) Water molecules resolved in TV map. Water molecules were only assigned to densities present in both maps generated using the two half datasets. Hydrogen bonds were shown in dashed lines with the bond length labeled.

In conclusion, by optimizing the cryo-EM grid preparation condition to plunge-freeze the immobilized particles on the antibody-affinity layer in vitreous ice with a thickness slightly larger than the particle size, we have obtained high-quality cryo-EM data for TV and solved a ~2.6 Å structure using the antibody-based affinity cryo-EM approach. This demonstrates the feasibility of using the antibody-based affinity cryo-EM approach for sub-3 Å structures and suggests the potential of using affinity cryo-EM approach as a more efficient and robust way of grid preparation for high-resolution cryo-EM. With antibodies commercially available for a vast number of samples, the antibody-based affinity cryo-EM method can potentially revolutionize cryo-EM sample preparation for a broader spectrum of native specimens, at least for most viral samples as demonstrated in this work. Optimization and demonstration of the antibody-based affinity cryo-EM approach for high resolution structure determination of smaller protein complexes will be performed in the near future for more general usage of this approach in cryo-EM.

## Materials and methods

### Purification of TV

TV was purified as described previously with some modifications (Yu et al., 2013). Briefly, LLC-MK2 cells (ATCC, Manassas, VA) grown to 80-90% confluency were inoculated with TV at a multiplicity of infection (MOI) of 0.2. Cells were collected after 72 hours and subject to three freeze-thaw cycles to release progeny TV. After clarifying by centrifugation at 13,776× g to remove cell debris, the supernatant was centrifuged at 28,000 rpm (SW28 rotor, Beckman, Danvers, MA). The pellet was suspended and further clarified by 20% (w/v) sucrose cushion ultracentrifugation at 28,000 rpm for 4 hours. Finally, the pellet was suspended and applied to a CsCl density gradient centrifugation at 288,000× g for 45 h (SW41Ti rotor, Beckman, Danvers, MA). The fractions containing TV virion were collected. Plaque assay was performed and revealed a titer of ~10^8^ plaque forming units (PFU)/ml. Sample was dialyzed against 1×PBS buffer (137 mM NaCl, 2.7 mM KCl, 10 mM phosphate, pH 7.4) for use.

### Preparation of antibody-coated TEM grids

The amorphous carbon layer of TEM grids was coated with antibodies as described before (Yu et al., 2016a, 2014). The protein A-aided antibody coating method was applied in this work. Briefly, homemade Formvar/carbon grids or commercial grids layered with lacey carbon and ultrathin continuous carbon (e.g., Ted Pella #01824) were glow-discharge cleaned for ~40 s at 15 mÅ current and 0.2-0.15 Torr vacuum using a Ladd vacuum evaporator (Ladd Research, Williston, VT). 5 μl of 50 μg/ml Protein A(Sigma Aldrich) was applied to the cleaned carbon surface and incubated for ~10-15 min at room temperature. After removing the excess liquids, 5 μl of diluted antiserum was quickly applied to the grid and incubated for 15-20 min in a humid chamber at ambient temperature. The residual liquid and unbound antibody were blotted away, and the grid was subjected to 30s washing with 100-200 μl of 1×PBS buffer (pH 7.4).

### Ice thickness estimation

Cryo-EM grids of frozen sample under different conditions were checked using a Philips CM200 microscope with a 1K CCD camera. The overall quality of the frozen grids were first visually evaluated as usual (Grassucci et al., 2007); For those promising conditions, the ice thickness was quantitatively estimated using the thickness estimation method based on Beer-Lambert law (Malis et al., 1988; Yan et al., 2015). More specifically, after melting the ice at a corner of the vitreous ice film, a low-magnification image (e.g., 3,800×) was taken covering both the region of interest and the melted area using a CCD camera. Beam intensities without (I_0_) and with (I) ice scattering, and inelastic mean free path of electrons in vitreous ice (λ_in_) determine the ice thickness: 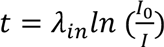. Since I_0_ and I can be measured directly using the DigitalMicrograph software (Gatan, Inc.) controlling the camera during sample screening, it provides immediate feedback for the ice thickness optimization. λ_in_ of 300 nm was used for the microscope operated at 200 KV as estimated in the references (Vulović et al., 2013; Yan et al., 2015).

### Affinity cryo-EM grid preparation and data collection of TV

TEM grids layered with lacey carbon support and a continuous ultrathin carbon (Ted Pella #01824) were coated with anti-TV antiserum (1:10 diluted, (Yu et al., 2014)) as described above. 3 μl of the purified TV sample was applied to the grid-coated with anti-TV antibody, and incubated in a humid chamber at 4 °C for ~30 min. Due to the limited amount of purified TV sample, only 3 μl sample was applied per grid. The grid was washed for ~2 min by floating on a drop of 200 μl cold 1×PBS, pH7.4, and plunge-frozen using a Gatan Cp3 (or FEI Vitrobot) cryo-plunger. Different cryo-EM grid freezing conditions (e.g., blotting time, humidity) were attempted and evaluated visually complemented by quantitative ice thickness estimation as aforementioned. A freezing condition leading to vitreous ice of ~60 nm on average for TV was identified. A dataset of ~1,000 movies was collected using FEI Titan Krios TEM operating at 300 KV and liquid nitrogen temperature, and recorded with the Gatan K2 Summit direct electron detector at super-resolution movie mode (5 frames/second) at 225,00× nominal magnification (0.65 Å/super-pixel). A dose rate of 8 electrons/physical pixel/second and a total exposure of 5 seconds resulted in a total dose of ~21 e/Å^2^ for the collected images.

### Single particle 3-D reconstruction using *jspr*

A script *motionCorrect.py* that runs the *dosefgpu_driftcorr* program (Li et al., 2013) in batch mode was used to determine motions among raw movie frames (https://github.com/jianglab/motioncorr). ~20,000 particles were selected from motion-corrected movie sums using RELION automatic particle picking module (Scheres, 2012). The *jspr* program *batchboxer.py* (Guo and Jiang, 2014; Liu et al., 2016) was used to extract the particles from raw movie frames based on the particle coordinates files, the frame shift parameters determined by *motionCorrect.py,* and the total electron dose. The extracted particles were both motion-corrected and dose-weighted adopting the radiation damage compensation strategy described in reference (Wang et al., 2014). The particle images were resampled from 0.650 to 0.975 Å/pixel using the Fourier cropping method. The *jspr* program *fitctf2.py* (Jiang et al., 2012) was used to determine the initial CTF parameters from the extracted particles. The estimation and correction of astigmatism aberration was performed later during iterative refinement using the aligner *refineAstigmatism* in *jspr* (Guo and Jiang, 2014; Liu et al., 2016). The whole dataset was then split into two half datasets, even and odd subsets, which were processed independently, including constructions of *de novo* initial models and subsequent refinements as described before (Guo and Jiang, 2014; Liu et al., 2016). Three correct *de novo* initial models were obtained using the random model method (Guo and Jiang, 2014; Liu et al., 2016) for both the “even” and “odd” halves, respectively, which were used as initial references to determine the Euler/Center parameters of all particles. This resulted in three sets of Euler/center parameters for each particle, based on which particles with consensus Euler/Center parameters were selected for further refinement. In the *jspr* program (Guo and Jiang, 2014), in addition to Euler/Center searching, many potential modulations present in 2-D images are also formulated as 2-D alignment tasks (oftenreferred to as high-order alignments), including micrograph and per-particle defocus, astigmatism aberration, overall scale variance among 2-D images, elliptic magnification distortion, and phase error from residual coma (Guo and Jiang, 2014; Liu et al., 2016; Yu et al., 2016b). Initial refinement was performed with only the Euler/Center parameters. After the convergence of the orientation/center refinement at ~3.6 Å, high-order aligners were gradually added to the iterative refinement loop to determine and correct those image artifacts as aforementioned, which all together improved the quality of the 3-D reconstruction to ~2.6 Å. The 0.143 cutoff for gold-standard Fourier Shell Correlation (FSC) curve (Rosenthal and Richard, 2003) was used to estimate the resolution of reconstruction with “even” and “odd” maps soft-masked using EMAN2 (Tang et al., 2007) adaptive masking processor (*mask.auto3d)*. The final map of TV was reconstructed from 14,154 particles by pooling the two half datasets. The final map was sharpened using the 1D structure factor of TV fitted from the images, and a low pass filter derived from the final FSC curve to boost the high-resolution Fourier amplitudes, and to suppress the high-frequency noise, respectively (Guo and Jiang, 2014; Rosenthal and Richard, 2003).

### Estimation of overall B factor (*B_overall_*) of cryo-EM data and reconstruction

Different numbers of particles were randomly selected from final refinement particle sets. New 3-D reconstructions were performed and the resolutions were estimated based on gold-standard FSC curves. The number of particles for 3-D reconstructions was then plotted against the corresponding achievable resolutions. Specifically, the *2ln(N_asu_d)* as a function of *1/d^2^* was plotted, in which *N_asu_* is the number of asymmetric units and *d*is the achievable resolution of 3-D reconstructions. Based on the mathematical formulation of the relationship of *N_asu_* d and the *B_overall_* (Figure 11 in (Rosenthal and Richard, 2003)), *B_overall_* can be approximated as the slope of the linear fitting of *2ln(N_asu_d)* and *1/d^2^*.

## Acknowledgements

This work was supported by NIH grant (1R01AI111095). Both negative staining and cryo-EM images were taken at Purdue Cryo-EM Facility. We thank Dr. Philip Serwer for providing the bacteriophage T7 antiserum, and Frank Vago for proofreading the manuscript.

## Supplementary Figures

**Figure s1.**
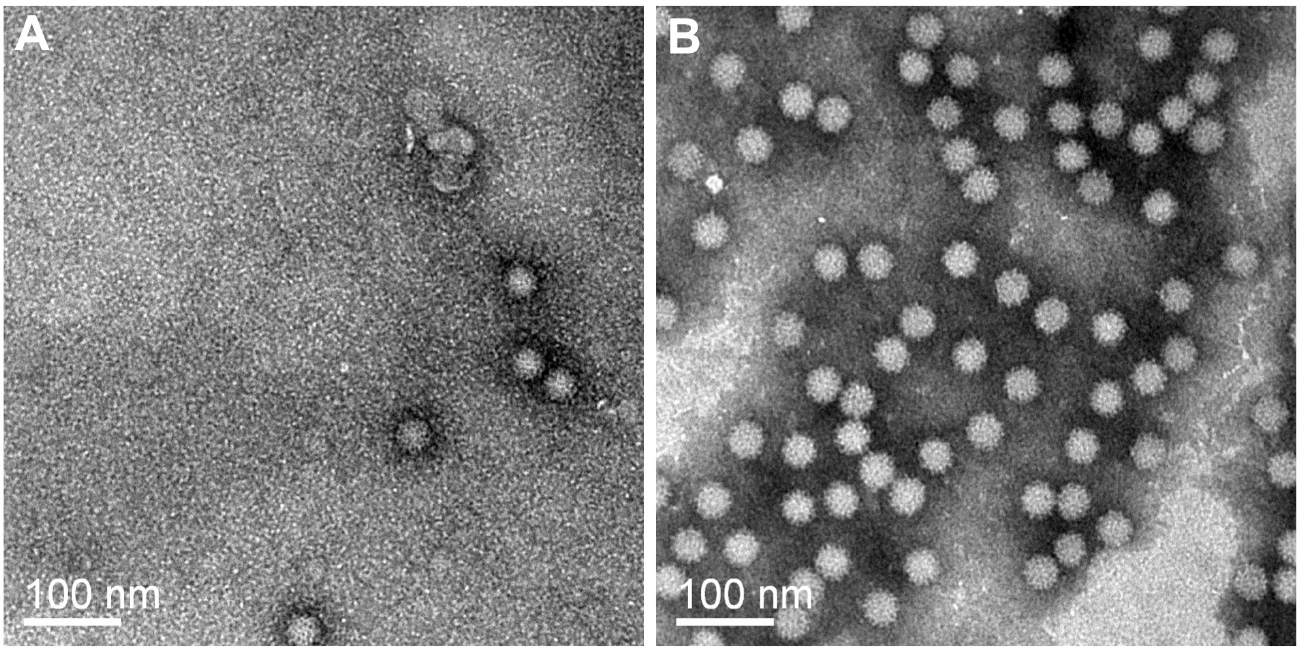
Negative stain TEM images of TV. (A) TV on regular TEM grid without affinity layer. (B) TV on the protein A-antiTV coated affinity grid. 3μl of purified TV sample was used to prepare both grids. Negative staining was performed with 2% phosphotungstic acid (PTA) at pH7.0.

**Figure s2.**
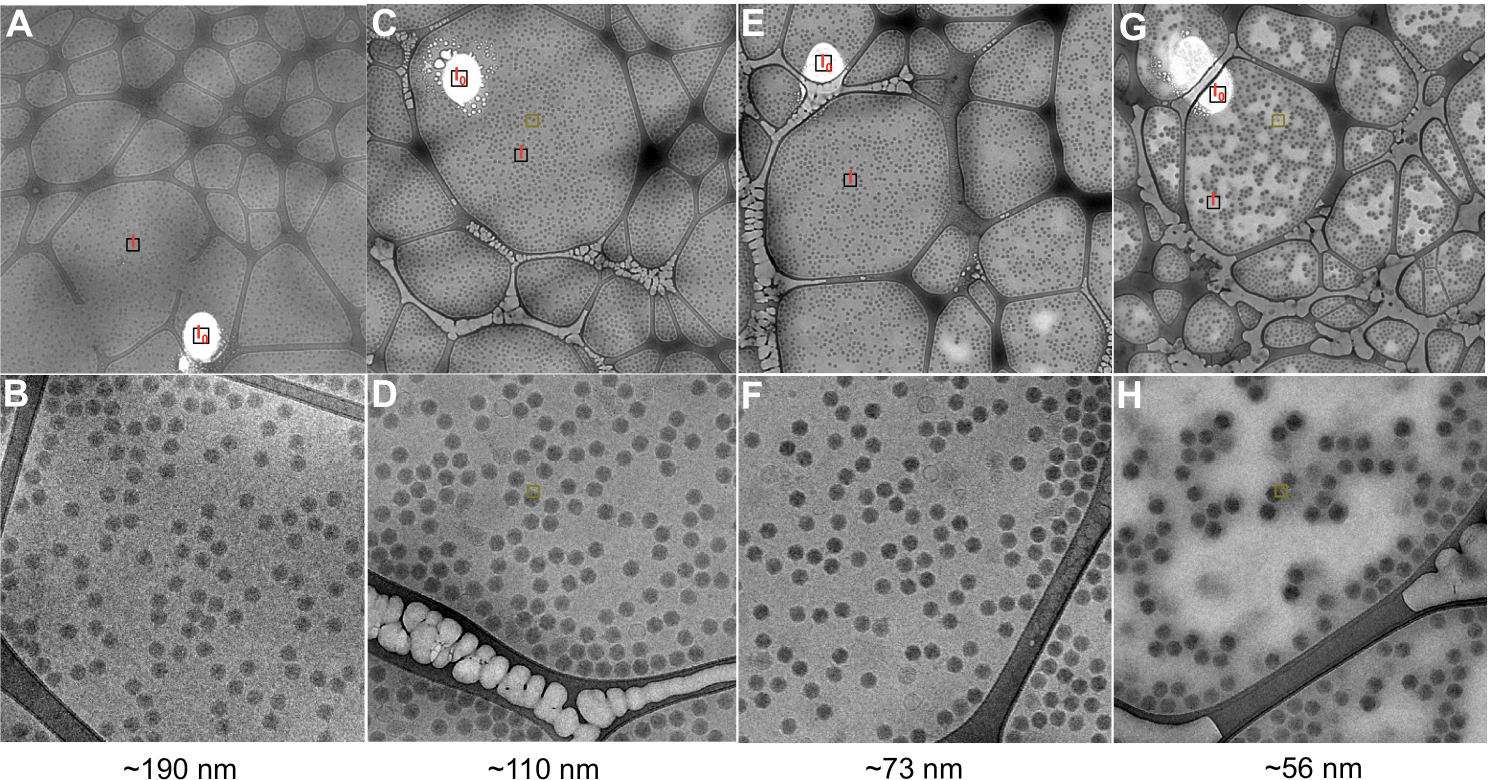
Optimization of cryo-freezing conditions of antibody affinity grids. Due to the limited amount of TV sample, the initial ice thickness optimization tests were performed using bacteriophage T7 that is available in large quantity in the lab. Affinity cryo-EM grids of bacteriophage T7 were prepared with protein A and anti-T7 antiserum as described in *Methods.*(A, C, E and G) show low magnification images of grids using four different freezing conditions. “I_0_” indicates the image intensities at the areas with ice melted away, and “I” represents intensities at areas with ice. The ice thickness of these four conditions was estimated at 190 nm, 110 nm, 73 nm and 56 nm, respectively, using the Beer-Lambert law based ice thickness estimation method. (B, D, F and G) show corresponding higher magnification micrographs of above four grids. Reduction of ice thickness correlated with the improvement of image contrast. The grid for micrograph G/H was slightly over-blotted that resulted in ice thickness of background areas (~56 nm) slightly smaller than the particle diameter (~70 nm).

**Figure s3.**
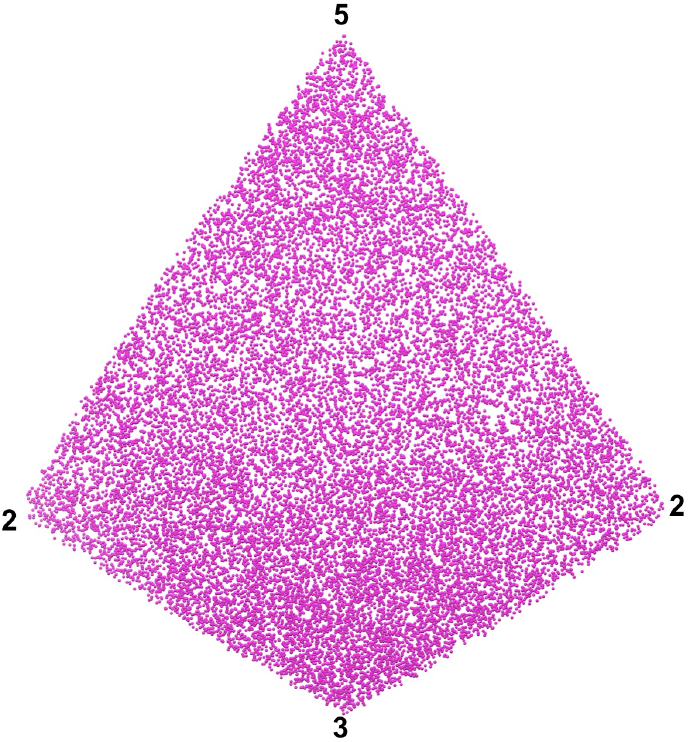
Euler angle distribution of TV particles captured by the antibody-based affinity cryo-EM grid. Figures “2”, “3” and “5” represent the icosahedral two-three-and five-fold axes, respectively.

**Figure s4.**
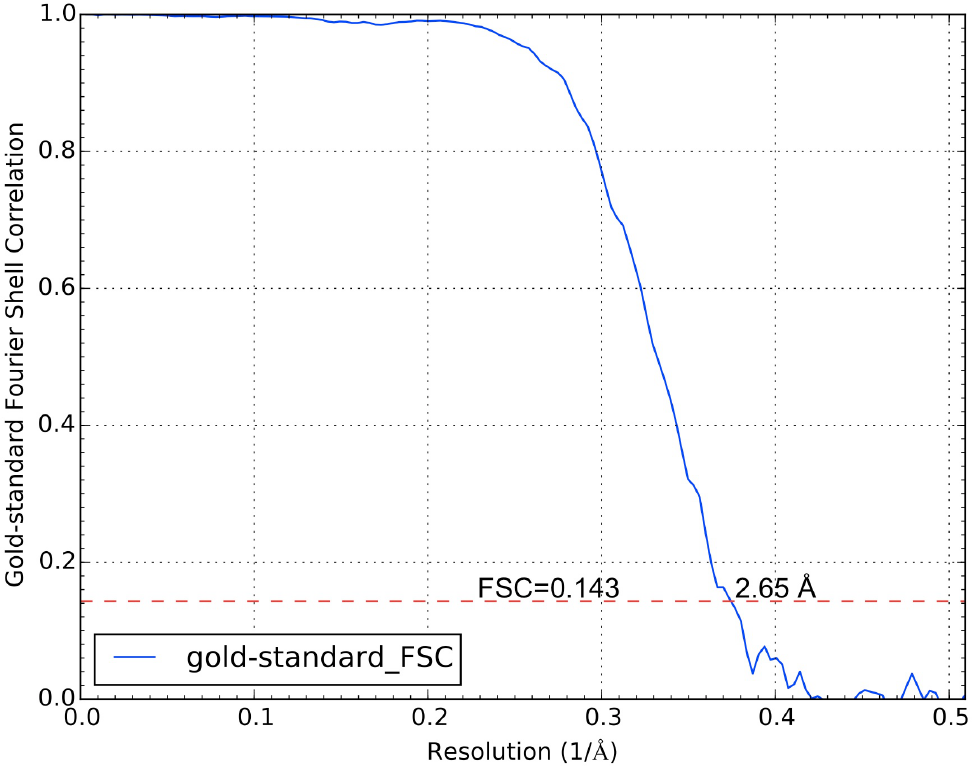
Gold-standard FSC curve of the 20S proteasome 3-D reconstruction with *jspr.* The resolution was estimated at 2.7 Å using the 0.143 cutoff. The dataset was downloaded from the public database EMPIAR (https://www.ebi.ac.uk/pdbe/emdb/empiar/entry/10025).

